## Extended data for "Multistability driven by cooperative growth in microbial communities"

**This PDF file includes:**

Extended Data Figs. 1 to 9

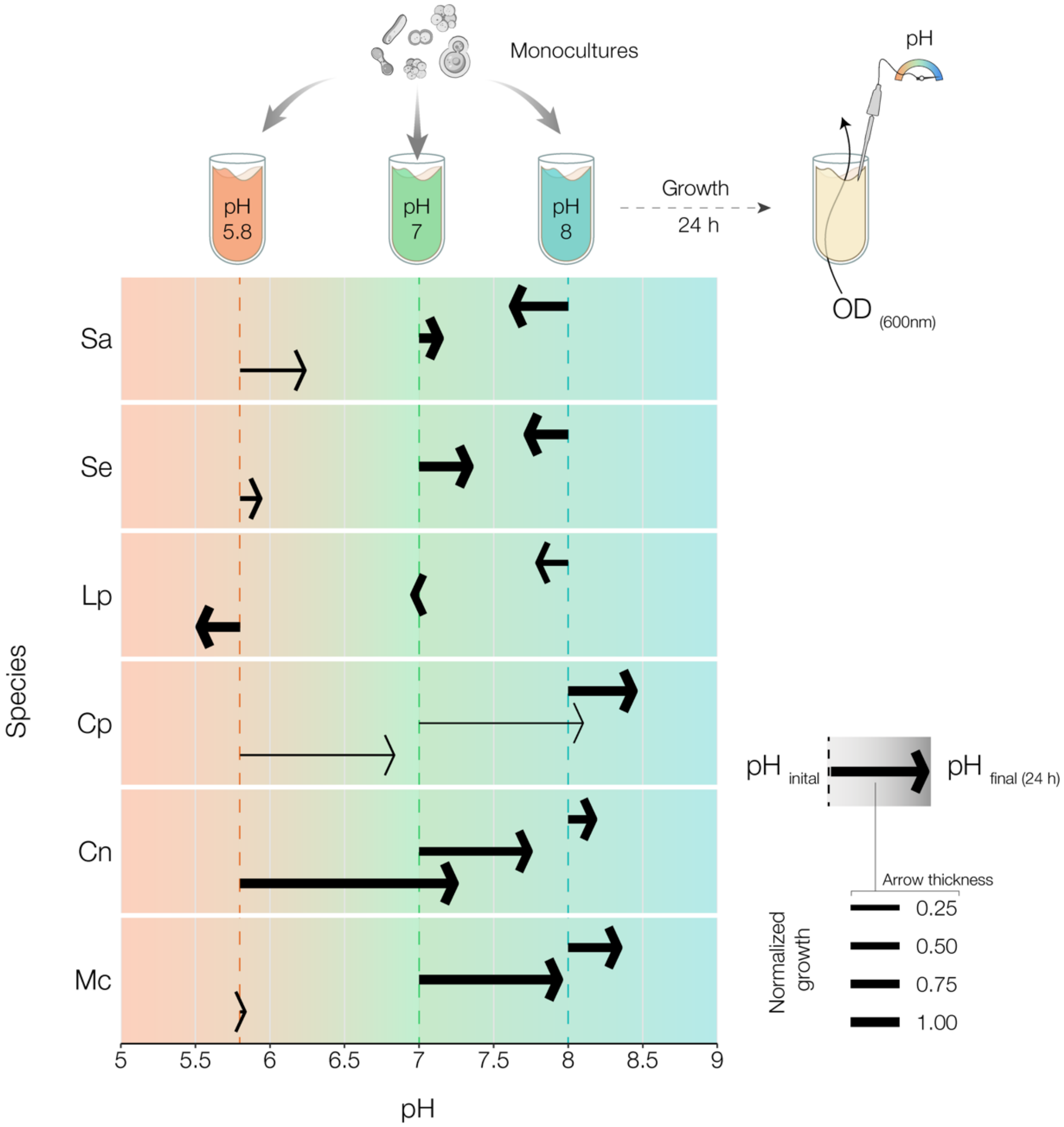


**Extended Data Fig. 1| The majority of the six species increase the pH of culture media independently of the initial pH.** Monocultures were grown in BHI at initial pHs: 5.8, 7 and 8. OD ($600_{nm}$) and pH were measured at the beginning of the experiment and after 24 h of growth. The length of arrows indicates pH changes induced by species at a given initial pH (average of four replicates). Arrow thickness is proportional to normalized growth and was calculated as the ratio between fold growth ($OD_{24h}/ OD_{0h}$) at a given initial pH and maximum species' fold growth across all initial pH conditions (average of four replicates). All focal species—*Cn*, *Mc*, *Sa* and *Se*—increase the pH of BHI media at initial pH 7, which was used in all the experiments described in the main text figures. *Lp* is the only species that lowers the pH in all conditions.

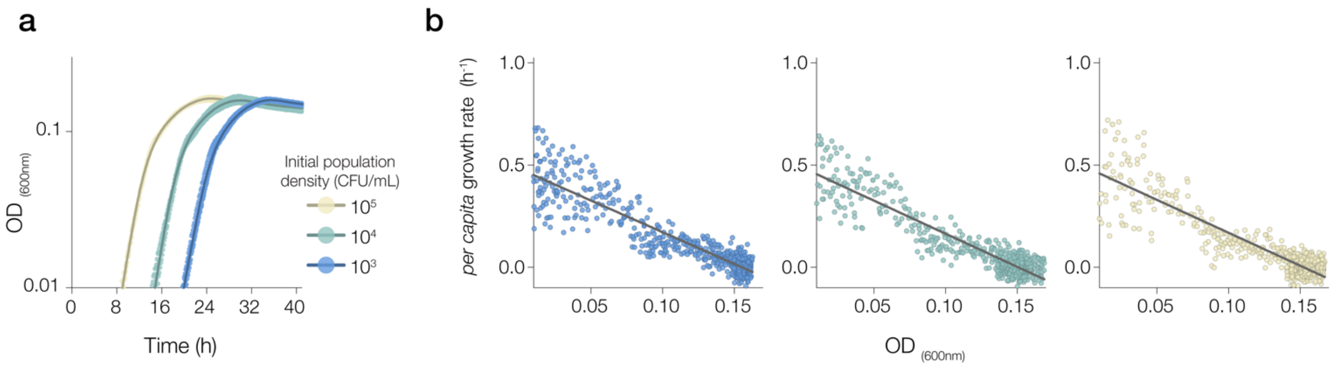


**Extended Data Fig. 2| *Cn* exhibits logistic growth.** **a,** To test whether the logistic growth model is a valid approximation for the growth of *Cn*, for which no Allee effect was experimentally observed (Figs. 2c and 4b), monocultures were grown at varying initial population densities over a period of 40 hours ($OD_{600nm}$ measurements every ~10 min, minimum of three replicates for each condition). **b,** In the logistic growth model, the *per capita* growth rate decreases linearly with the population density. The data show the local *per capita* growth rate calculated as $\ln OD_{t+\Delta t} - \ln OD_t \,/\, \Delta t$, with the interval between measurements $\Delta t$=10min, as a function of the population density ($OD_{600nm}$). Lines show the results of a linear fit for populations initiated at $10^3$ CFU/mL (slope = -3.105, $R^2$ = 0.8372), $10^4$ CFU/mL (slope = -3.240, $R^2$ = 0.8452) and $10^5$ CFU/mL (slope = -3.224, $R^2$ = 0.8059). For three different initial population densities, the data reveal an approximately linear decrease of the *per capita* growth rate with the population density, which confirms that *Cn* populations grow logistically.

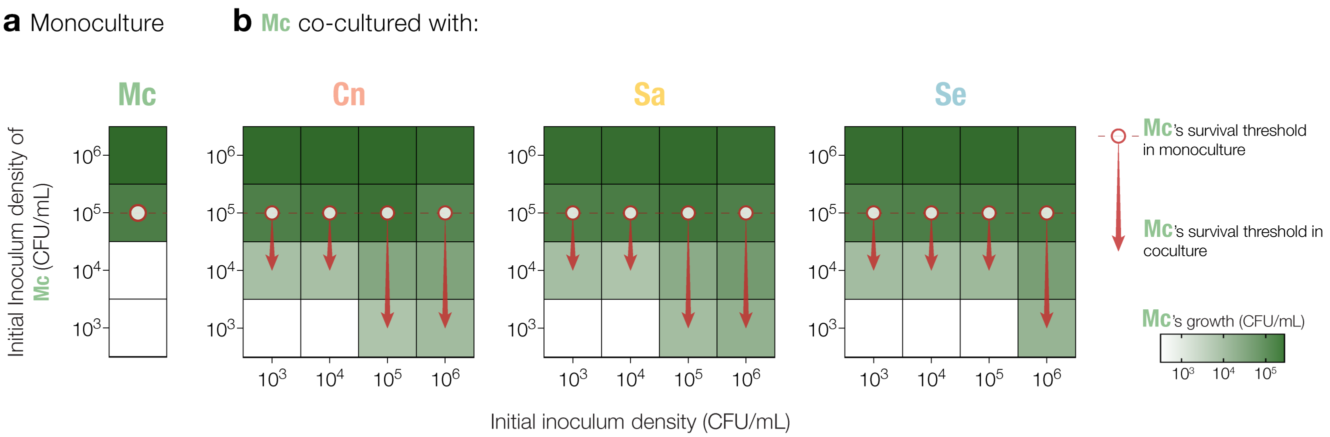


**Extended Data Fig. 3| *Mc*'s survival threshold is lower in coculture with all competitors. a,** *Mc* monoculture inoculated in BHI at varying initial inoculum densities. CFU counting was performed after 6 h of incubation to determine population density (CFU/mL). No growth is observed below the survival threshold of *Mc* (~$10^5$ CFU/mL, white dot). **b,** *Mc* co-cultured with varying initial inoculum densities of *Cn*, *Sa* or *Se*. These three species facilitate the growth of *Mc*, lowering its survival threshold (arrow) in comparison to growth in monoculture. Data obtained from the average of 3 replicates for each condition.

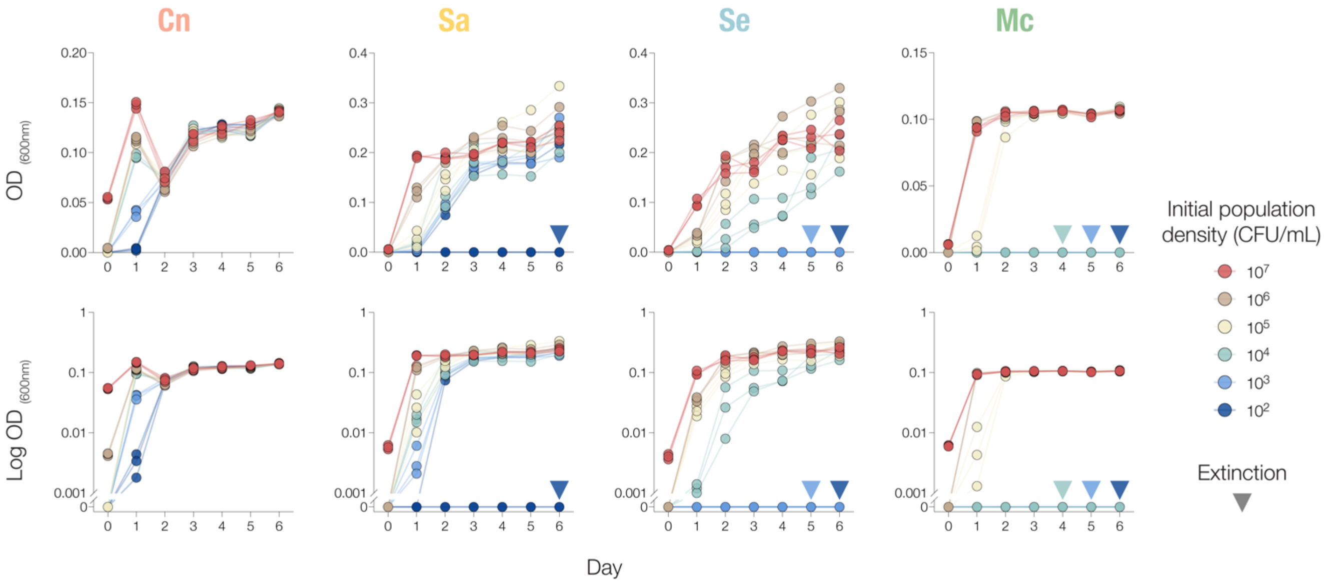


**Extended Data Fig. 4| Population dynamics of species with the Allee effects exhibit survival thresholds under daily dilutions.** We propagated monocultures at varying initial population densities over six days using a dilution factor of 100X. OD measurements are shown on linear (top) and log scales (bottom) for all focal species (n=3). Species subject to the Allee effect—*Sa*, *Se* and *Mc*—go extinct (inverted pyramid) when inoculated below critical (threshold) density. Replicate communities starting near the critical density may split, with some populations surviving and others going extinct.

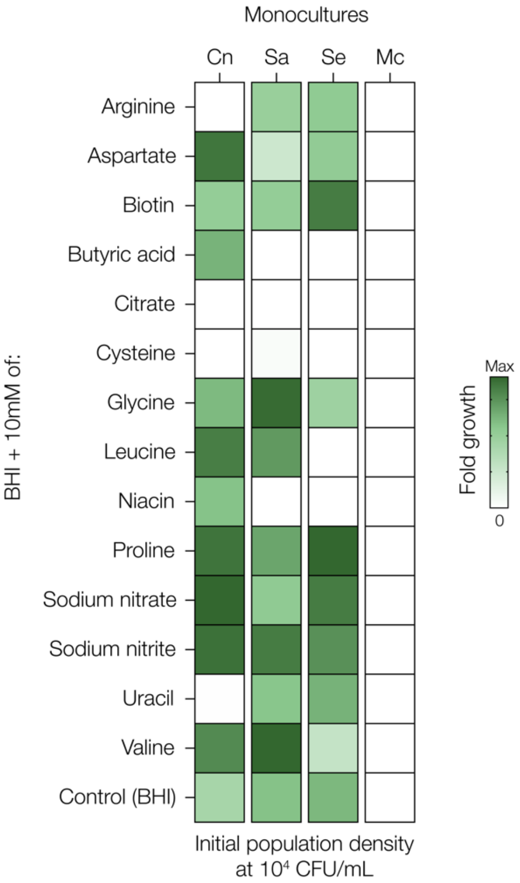


**Extended Data Fig. 5| Biochemical screening reveals molecules capable of promoting species growth at initially low population density.** We selected a set of molecules important to central metabolic pathways to supplement BHI (see Fig. 4a for additional molecules). Heatmap shows the monoculture fold growth ($OD_{24h}/OD_{0h}$, n=3) of focal species inoculated at low population density ($10^4$ CFU/mL) in supplemented BHI and control (BHI only, bottom). Maximum fold growth is scaled for each column relative to the maximum value observed for the corresponding species across conditions. *Mc*'s growth below survival threshold (~$10^5$ CFU/mL) is only observed in glutamate supplementation (Fig. 4a).

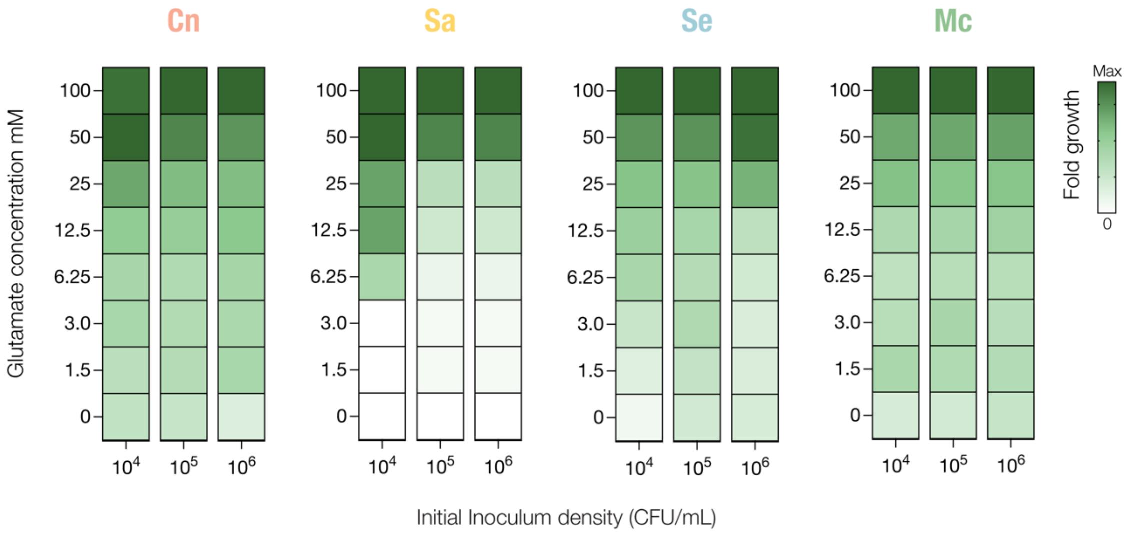


**Extended Data Fig. 6| Dose response assay of increasing concentrations of glutamate shows maximum fold growth of species with the Allee effects at a concentration of 100mM.** Heatmap shows the fold growth ($OD_{24h}/OD_{0h}$, n=3) of focal species in BHI supplemented with increasing concentrations of glutamate. Maximum fold growth occurred at a concentration of 100mM for *Sa*, *Se* and *Mc*. Maximum fold growth is scaled for each column according to initial inoculum density.

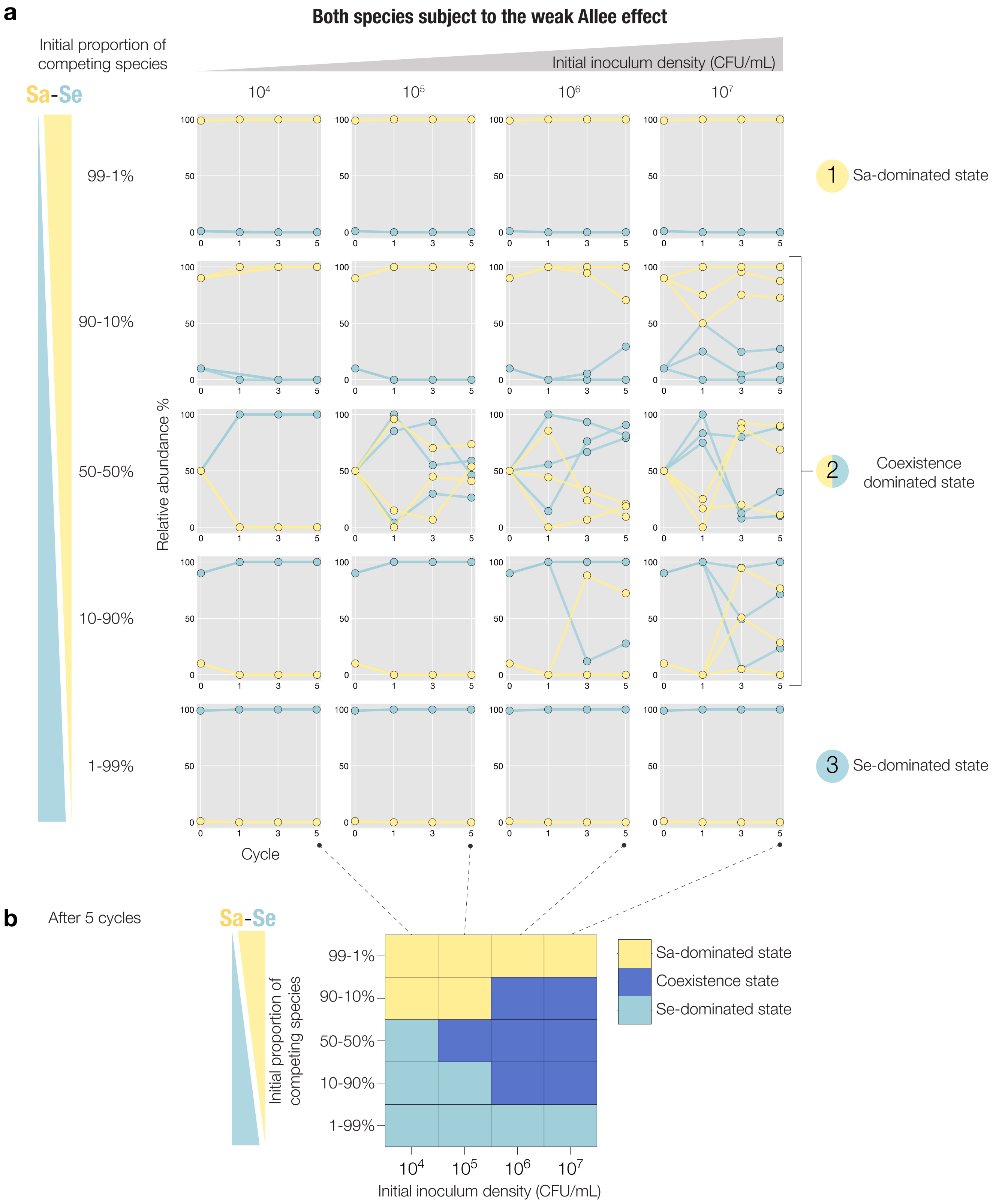


**Extended Data Fig. 7| Tristability emerges in pairwise competition between Sa and Se, which exhibits both weak interspecies interactions and weak Allee effect.** **a,** Results of additional experiments characterizing the competitions between *Sa* and *Se* mapped across various initial inoculum densities and different initial proportions of competing species. The non-canonical outcome of tristability is observed after 5 growth-dilution cycles, with initial inoculum densities ranging from $10^5$ to $10^7$ CFU/mL. The tristable outcome is composed of two states analogous to those of bistability (*Sa*-dominated state *1* and *Se*-dominated state *3*) and coexistence between species (state *2*). Time trajectories correspond to three replicates for each condition. Variability in replicates of the coexistence state indicates that tristability might be less stable. **b,** Phase diagram mapping the three stable states on the space of inoculum densities (average of three replicates shown in panel a).

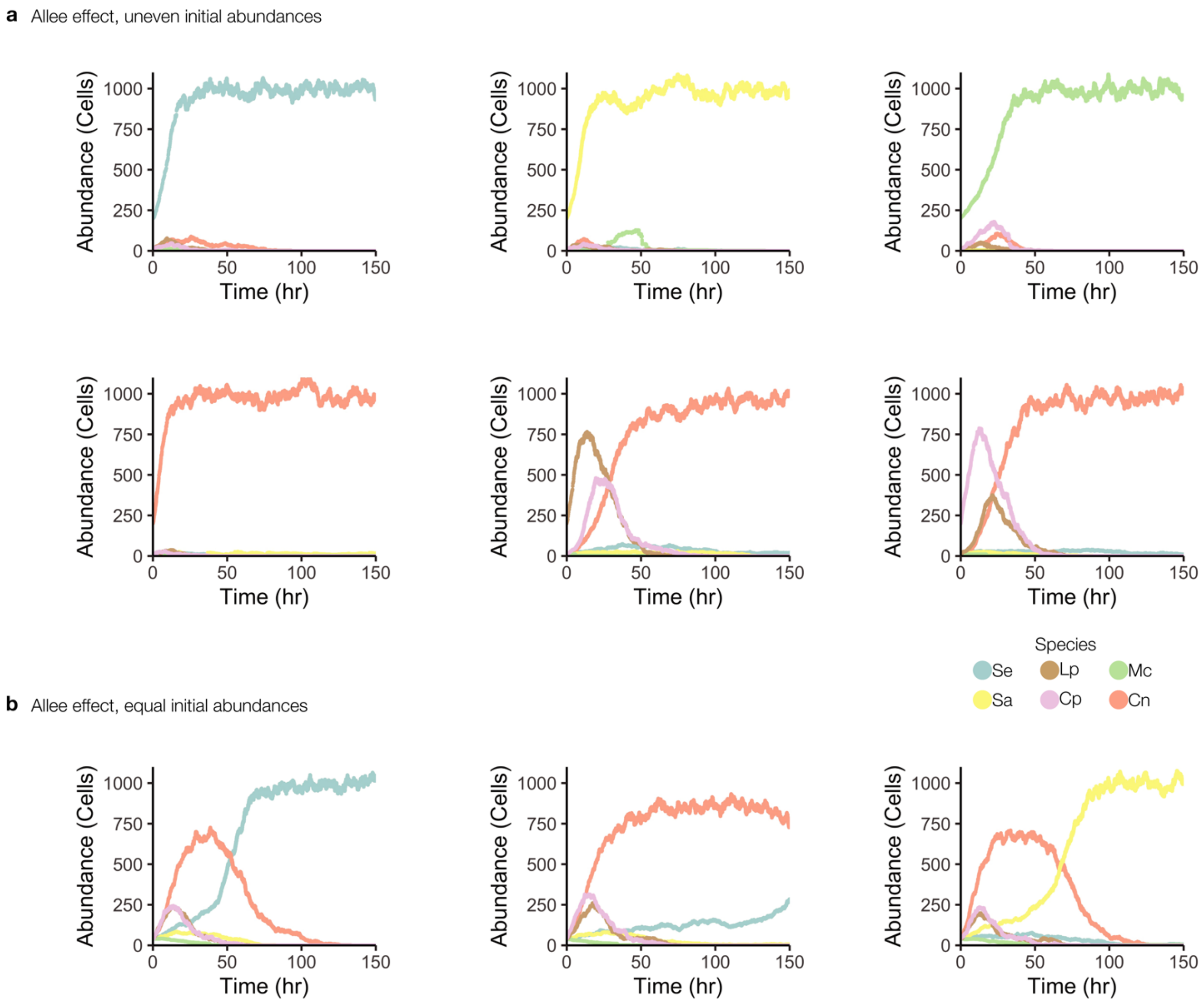


**Extended Data Fig. 8| Sensitivity to stochastic fluctuations can lead to divergent community outcomes from the same initial conditions. a,** Representative time series from stochastic simulations in which communities are initiated at uneven species abundances where one species is initially dominant. For a fixed set of parameter values (Extended Data Text 1) the stochastic model recapitulates the 4 stable states observed experimentally. **b,** Starting from equal abundances for all six species, stochastic fluctuations can lead the community to reach alternative outcomes. The three panels show representative time series in which the community reaches the stable state dominated by *Se* (left), a case of slow dynamics in which *Cn* still coexists with *Se* (center) after 150 simulated hours (~6 days), and the reaching of the stable state dominated by *Se* (right). These three representative outcomes of stochastic simulations recapitulate the observed outcomes for communities inoculated at equal initial abundances (Fig. 1c), suggesting that sensitivity to stochastic fluctuations can lead the community to reach different outcomes from identical, or similar, initial conditions.

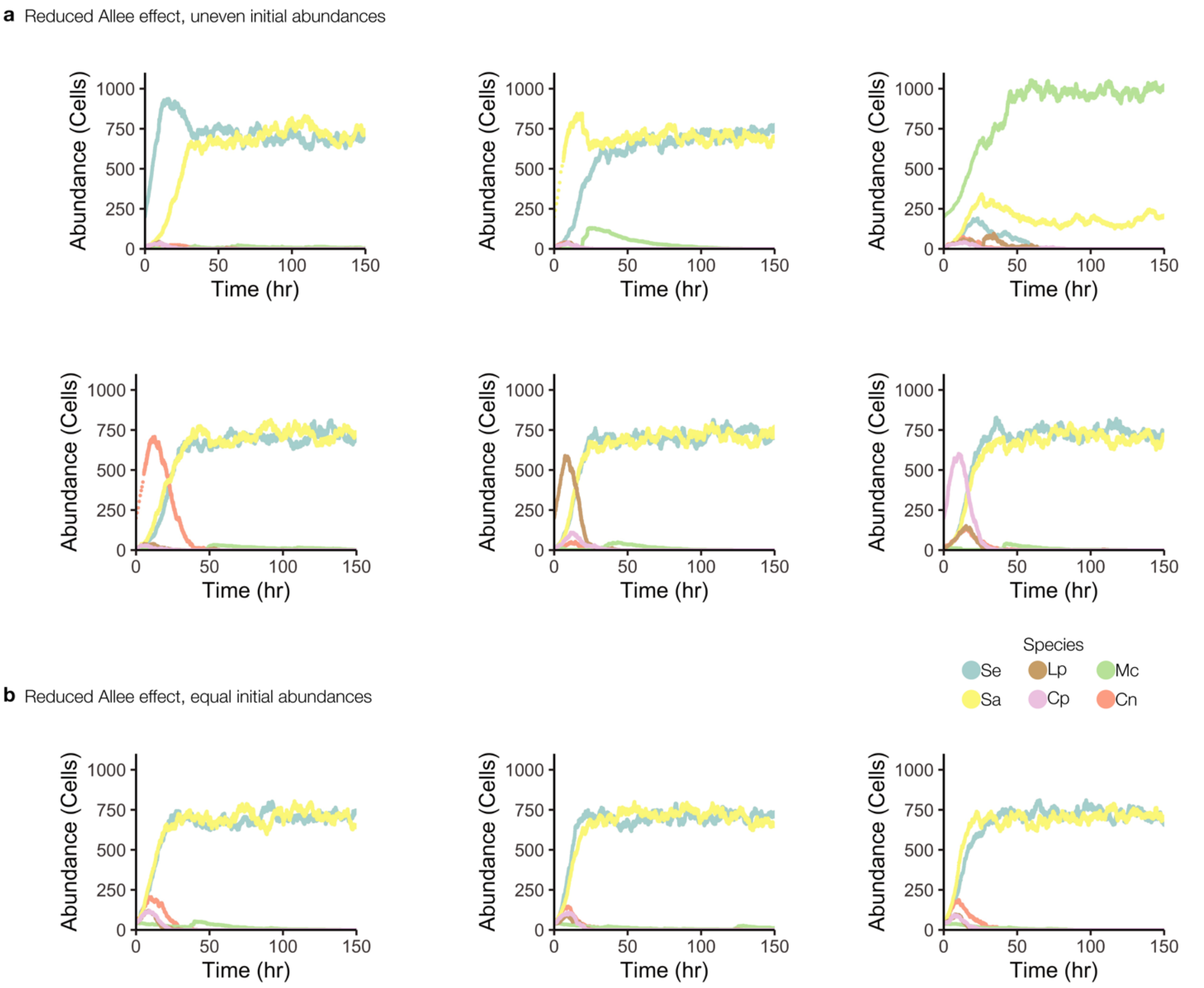


**Extended Data Fig. 9| Reducing the Allee effect in stochastic populations recapitulates the observed changes in the number of alternative stable states.** **a,** Representative time series from stochastic simulations in which communities are initiated at uneven species abundances for a reduced Allee effect scenario, which corresponds to the case of experimental cocultures under glutamate supplementation. Simulations recapitulate the reshaping of alternative stable states associated with the reduced Allee effect: the community reaches a stable state in which *Se* and *Sa* coexist under a wide range of different initial conditions. For the specific case in which *Mc* begins at relatively high density, the community reaches an alternative stable state in which *Mc* and *Sa* coexist. In these simulated scenarios, all the species grow logistically (Methods) except *Mc*, which is subject to the strong Allee effect (parameter value of the Allee threshold set to $c=0.02$). Parameter values are otherwise identical to those in Extended Data Fig. 8 (Supplementary information Text 1). **b,** Starting from equal abundances for all six species, stochastic fluctuations under the reduced Allee effect preferentially reach the stable state dominated by *Se* and *Sa* (this scenario is shown for direct comparison with Extended Data Fig. 8 although it was not experimentally tested).
