## Supplementary information for "Multistability driven by cooperative growth in microbial communities"

**This PDF file includes:**

Extended Data Text 1

**Extended Data Text 1 | Stochastic model and parameter values.** To apply the optimized tau-leap method in the stochastic simulations (Extended Data Figs. 8 and 9), we used the equations of the theoretical model (Methods) in their non-normalized version. In the absence of an Allee effect, the dynamics of a given species *i* therefore reads:

$$\frac{dx_i}{dt} = r_i x_i\left(1 - \alpha_{ij} x_j / K_j\right),$$

Where $x_i$ stands for the absolute (non-normalized) abundance of species i, $K_i$ for its carrying capacity, and the rest of the terms are analogous to those in the normalized version of the model (Methods).

For species that are subject to an Allee effect, the non-normalized version of the dynamics reads:

$$\frac{dx_i}{dt} = r_i x_i\left((x_i/K_i - a_i)(1 - x_i/K_i) - \alpha_{ij} x_j / K_j\right),$$

Under the convention *i=1,2,3,4,5* and *6* corresponding to Se, Cn, Sa, Mc, Lp and Cp, respectively, we used the following parameter values:

$$r_i = 0.3\ hr^{-1}, \text{for } i = [1,6],$$

$$\boldsymbol{\alpha} = \begin{pmatrix} 1 & 0.4 & 0.4 & 1.2 & 0.1 & 0.1 \\ 1.05 & 1 & 1.2 & 1.6 & 0.3 & 0.3 \\ 0.4 & 0.4 & 1 & 0.8 & 0.1 & 0.1 \\ 0.1 & 0.1 & 0.1 & 1 & 0.1 & 0.1 \\ 1.3 & 1.3 & 1.3 & 1.5 & 1 & 0.1 \\ 1.3 & 1.3 & 1.3 & 1.5 & 0.1 & 1 \end{pmatrix}$$

In the simulations in Extended Data Fig. 8, we used the Allee effect strength values:

$$\boldsymbol{c} = (c_1, c_3, c_4) = (-0.3, -0.27, 0.05)$$

To model the reduction of the Allee effect under glutamate supplementation, in the simulations in Extended Data Fig. 9, only *Mc* (*i=4*) was subject to the Allee effect (with a reduced strength $c_4$=0.02).

It is worth noting that the parameter values used in the main text (Figs. 3b, 5a and 5b) were chosen in order to favor the visualization, in the corresponding phase planes, of the qualitative impact of the Allee effect on pairwise competitions outcomes. While the parameter values in the stochastic simulations (Extended Data Figs. 8 and 9) generate qualitatively the same pairwise outcomes presented in the main text, the set of parameter values used for the stochastic simulations enables a better recapitulation of the overall experimental outcomes, including the divergent outcomes for the 6-species community (Extended Data Fig 8b).
